# Candidate urine biomarker discovery from only five pairs of samples before and after tumor resection in glioma patients

**DOI:** 10.1101/240861

**Authors:** Jianqiang Wu, Jun Zhang, Yuanli Zhao, Youhe Gao

**Author notes:** JW and JZ contributed equally to this work. **Corresponding author**: Prof. Zhao Yuanli; Prof. Youhe Gao.

## Abstract

Biomarkers are measurable changes associated with the disease. Without the control of homeostatic mechanisms, urine accumulates systemic body changes and thus serves as an excellent early biomarker source. However, urine is affected by many factors other than disease. Although many candidate biomarkers have been identified in animal models, a large number of clinical samples might still be required for the disease related changes. A self-controlled study should be able to avoid the interferences of individual differences among patients. Gliomas are the most common primary malignant brain tumors and have a very poor prognosis. Early diagnosis of gliomas and the monitoring of tumor recurrence are crucial to improve glioma patient outcomes. Here we set to try if biomarker candidates can be identified by comparing urine samples from five glioma patients collected at the time of tumor diagnosis and after surgical removal of the tumor. Using label-free liquid chromatography coupled with tandem mass spectrometry (LC-MS/MS) quantification, twenty-seven urinary proteins were significantly changed after tumor resection (fold change ≥ 1.5, P-value < 0.05, and similar changes in all 5 patients), many of which have been previously associated with gliomas, such as CEACAM1, ANXA7, CALR, CRYAB, CD276, pIgR and cathepsin D. Functions of these proteins were significantly enriched in the regulation of tissue remodeling, autophagy, the inhibition of gene expression, the positive regulation of natural killer cell-mediated cytotoxicity and angiogenesis, which are associated with glioma development. Our results suggested that using the self-control of before and after tumor resection is an effective method to identify differential proteins associated with the disease, even with a small number of patients.

## Introduction

Gliomas are the most common primary brain tumor in adults, representing 81% of malignant brain tumors [1]. Malignant gliomas have a very poor prognosis and a high risk of tumor recurrence after treatment. Therefore, timely diagnosis of gliomas and glioma patient monitoring for disease recurrence are essential to improve patient survival. There are no currently generally accepted screening protocols to reveal asymptomatic brain tumors [2]. Thus, there is an urgent need to discover new biomarkers for noninvasive diagnosis and to monitor tumor recurrence in glioma patients.

Urine is a sensitive biofluid for biomarker research. Without the control of homeostatic mechanisms, urine accumulates systemic bodily changes and thus is an excellent biomarker source [3]. A roadmap for the urine biomarker era was previously proposed [4], and many candidate urine biomarkers were successfully identified in animal models [5–15]. In these animal models, the effects of genetic and environmental factors on the urine proteome were limited to the minimum, which is useful when identifying disease biomarkers. However, patient urine can be affected by many factors, including gender, age, exercise, and hormone conditions [16]; thus, many clinical samples are required to identify verifiable disease biomarkers. A self-controlled study should be able to avoid the interferences of individual differences among patients. It is unknown whether promising biomarkers can be identified from the relatively smaller number of glioma patient urine samples before and after tumor resection.

In this study, urine samples from five glioma patients were collected at the time of tumor diagnosis and after total surgical removal of brain tumors. Postoperative urine samples were collected 1–2 weeks after surgery and just before discharge to avoid trauma effects from altering the urine proteome. A comparative proteomic analysis of urine samples before and after tumor resection was performed using liquid chromatography coupled to tandem mass spectrometry (LC-MS/MS).

## Materials and Methods

### 1. Patients

Patients were recruited from the Peking University International Hospital. All patients had evident tumors identified in magnetic resonance imaging (MRI) studies at preoperative urine collection. The final tumor diagnosis was based on the histopathology of resected tissues. None of the cancer patients were treated with chemotherapeutic agents or radiation during sample collection. None of the adult patients had known histories of vascular malformation or recent surgery. The gross total resection of brain tumors was also documented by imaging.

All subjects were informed about the purpose of the study and signed informed consent forms before study inclusion. The study was approved by the local ethical committee of Peking University International Hospital.

### 2. Urine collection and sample preparation

Midstream first morning urine was collected from five glioma patients before and after tumor resection. The preoperative urine samples were collected from patients one day before surgery. The postoperative samples were collected 1–2 weeks after the patient recovered from surgery to avoid effects from stress. After collection, urine samples were centrifuged at 5,000 g for 30 min at 4°C to remove cell debris and contaminants. The supernatant was precipitated with three volumes of ethanol at −20°C overnight followed by centrifugation at 12,000 g for 30 min. The pellet was resuspended in lysis buffer (8 M urea, 2 M thiourea, 50 mM Tris, and 25 mM DTT). The protein concentration of each sample was measured using the Bradford assay.

### 3. Tryptic digestion

Urinary proteins were prepared using the FASP method. Protein samples (100 µg) were denatured with 20 mM dithiothreitol at 37°C for 1 h and alkylated with 50 mM iodoacetamide in the dark for 30 min. The samples were loaded onto 10 kD filter devices (Pall, Port Washington, NY, USA) and centrifuged at 14,000 g and 18°C. After washing twice with UA (8 M urea in 0.1 M Tris-HCl, pH 8.5) and four times with 25 mM NH_4_HCO_3_, the samples were digested with trypsin (enzyme-to-protein ratio of 1:50) at 37°C overnight. The peptide mixtures were desalted using Oasis HLB cartridges (Waters, Milford, MA) and dried by vacuum evaporation.

### 4. LC-MS/MS analysis

Ten peptide samples resulting from the above digestion were re-dissolved in 0.1% formic acid to 0.5 µg/µL. For analysis, 1 µg of peptides from an individual sample was loaded onto a trap column and separated on a reverse-phase C18 column (75 µm × 100 mm, 2 µm) using the EASY-nLC 1200 HPLC system (Thermo Fisher Scientific, Waltham, MA, USA). The analytical column was eluted over 60 min at a flow rate of 300 nL/min. The peptides were analyzed with an Orbitrap Fusion Lumos Tribrid mass spectrometer (Thermo Fisher Scientific, Waltham, MA, USA). MS data were acquired in high-sensitivity mode using the following parameters: data-dependent MS/MS scans per full scan with top-speed mode (3 s), MS scans at a resolution of 120,000, MS/MS scans at a resolution of 30,000 in Orbitrap, 30% HCD collision energy, charge-state screening (+2 to +7), dynamic exclusion (exclusion duration 30 s), and a 45 ms maximum injection time.

To identify more urinary proteins and validate the differential proteins identified above, the ten peptide samples were reanalyzed using a 90 min elution time, keeping the other MS parameters the same as above.

### 5. Label-free proteome quantification

The proteomic data were searched against the SwissProt Human database (released in 2017 and containing 20,169 entries) using Mascot software (version 2.6.1, Matrix Science, London, UK). The parent ion tolerance was set to 10 ppm, and the fragment ion mass tolerance was set to 0.05 Da. Carbamidomethyl cysteine was set as a fixed modification, and the oxidation of methionine was considered a variable modification. The specificity of trypsin digestion was set for cleavage after K or R, and two missed trypsin cleavage sites were allowed. Peptide and protein identification was further validated using Scaffold software (version 4.8.4, Proteome Software Inc., Portland, OR). Peptide identifications were accepted at an FDR less than 1.0% using the Scaffold Local FDR algorithm and with at least 2 unique peptides. Comparisons across different samples were performed after normalization to total spectra accounts using Scaffold software. Spectral counting was used to compare the protein abundance between two groups according to a previously described procedure [17, 18].

### 6. Statistical analysis

Statistical analysis was performed with SPSS. Comparisons between the two groups were conducted using paired Student’s t-test. Group differences resulting in P-values < 0.05 were considered statistically significant.

## Results

### 1. Clinical characteristics of glioma patients

A total of 5 consecutive and histopathology diagnosed glioma patients were recruited from Peking University International Hospital between April 2017 and May 2017. Their ages, sexes, tumor locations, histology result and degree of resection are summarized in **Table 1**. None of them were receiving any chemotherapeutic agents or radiation during sample collection and had no histories of vascular malformations or recent surgery.

**Table 1.** Clinical characterstics of the five glioma patients.

| Patient No. | Sex (M/F) | Age (yrs) | Tumor location | Histology <sup>a</sup> | Degree of resection <sup>b</sup> |
| --- | --- | --- | --- | --- | --- |
| 1 | M | 39 | Right temporal lobe | Grade IV | 100% |
| 2 | M | 49 | Right temporal lobe | Grade IV | 100% |
| 3 | F | 39 | Right frontal lobe | Grade IV | 100% |
| 4 | F | 62 | Left temporal lobe | Grade IV | 100% |
| 5 | F | 37 | Brainstem | Grade IV | 90% |

**Note:** <sup>a</sup> glioma grade classified as World Health Organization (WHO) grades I–IV; <sup>b</sup> according to the volume of tumor before and after operation measured on the contrast-enhanced MRI.

All of five patients had malignant glioma (WHO grade IV) convinced by the histopathology of resected tissues. To measure the degree of resection, all of them had contrast-enhanced MRI scanning before and after tumor resection. An image of a representative patient brain before and after tumor resection is shown in **Figure 1**.

**Figure 1.**
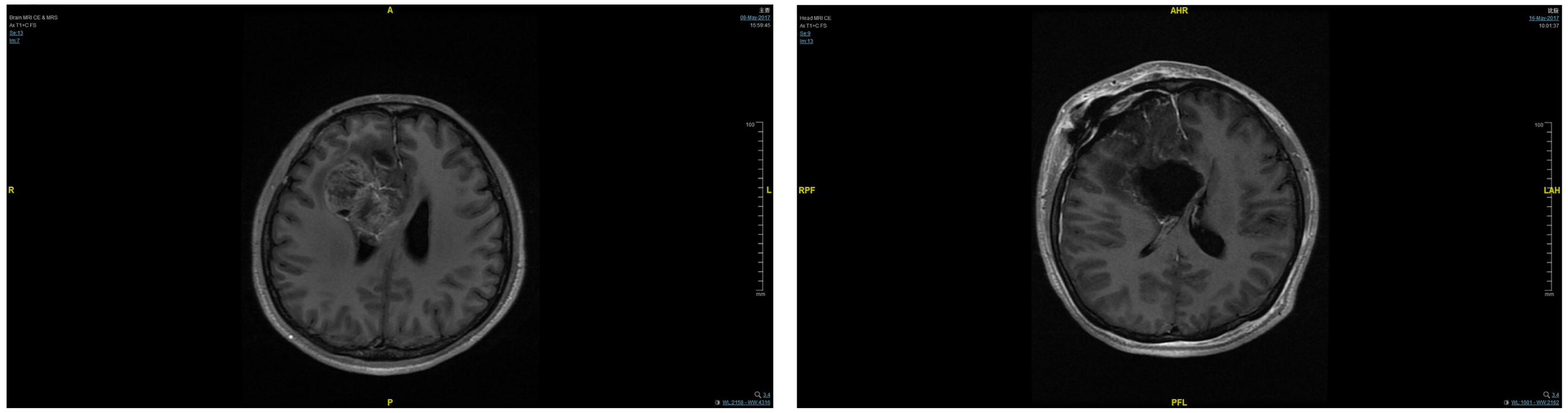
MRI of brain before and after tumor resection.

### 2. Changes to the urine proteome after tumor resection

Ten urine samples from five glioma patients before and after tumor resection were collected, and label-free LC-MS/MS quantification was used to characterize the differential expression of urinary proteins. A total of 1,377 urinary proteins with at least two unique peptides were identified with < 1% FDR at the protein level. All identification and quantification details are presented in **Table S1**.

From the spectral counts of the identified proteins, there were significant individual differences between glioma patient samples. Urine samples were divided into a patient group and treatment group, which indicated the condition of glioma patients before or after tumor resection. Differential proteins were screened with the following criteria: fold change ≥ 1.5 between two groups and P-value of paired t-test < 0.05. All five patient samples indicated similar changes in protein level after surgical treatment. A total of 27 differential proteins were identified; their details are described in **Table 2**. Of these 27 proteins, the eight most significant were Carcinoembryonic antigen-related cell adhesion molecule 1 (CEACAM1), mitochondrial 10-kDa heat shock protein (HSPE1), adhesion G protein-coupled receptor F5 (ADGRF5), mitochondrial malate dehydrogenase (MDH2), intestinal maltase-glucoamylase (MGAM), acid ceramidase (ASAH1), calreticulin (CALR), and thrombospondin-4 (THBS4) (**Figure 2**).

**Table 2.** Differential proteins identified in urine samples before and after tumor resection in glioma patients.

| Uniprot | Description | FC <sup>a</sup> | P | Glioma tissue <sup>b</sup> | Related to gliomas <sup>c</sup> |
| --- | --- | --- | --- | --- | --- |
| **P13688** | Carcinoembryonic antigen-related cell adhesion molecule 1<br>(CEACAM1) | ↑∞ | 0.052 |  | [20] |
| P27797 | Calreticulin (CALR) | ↓∞ | 0.068 | Y | [24] |
| **P35443** | Thrombospondin-4 | ↓13 | 0.024 |  |  |
| **P61604** | 10 kDa heat shock protein, mitochondrial | ↑8.5 | 0.018 | Y |  |
| **Q8IZF2** | Adhesion G protein-coupled receptor F5 | ↑7 | 0.018 |  |  |
| P20073 | Annexin A7 (ANXA7) | ↑6 | 0.034 | Y | [21] |
| **P40926** | Malate dehydrogenase, mitochondrial | ↑4.17 | 0.007 | Y |  |
| P02511 | Alpha-crystallin B chain (CRYAB) | ↑3.90 | 0.033 | Y | [26] |
| P09417 | Dihydropteridine reductase | ↑2.5 | 0.051 | Y | [12] |
| **O43451** | Maltase-glucoamylase, intestinal | ↑2.43 | 0.010 | Y |  |
| **P13489** | Ribonuclease inhibitor | ↓2.4 | 0.049 | Y |  |
| **Q9H3G5** | Probable serine carboxypeptidase CPVL | ↑2.31 | 0.015 | Y |  |
| **P13473** | Lysosome-associated membrane glycoprotein 2 | ↑2.15 | 0.012 | Y | [34] |
| Q92820 | Gamma-glutamyl hydrolase | ↑2.02 | 0.015 | Y |  |
| **Q07075** | Glutamyl aminopeptidase | ↑2.00 | 0.046 | Y | [12] |
| **Q5ZPR3** | CD276 antigen (CD276) | ↑1.92 | 0.042 | Y | [29] |
| O96009 | Napsin-A | ↑1.90 | 0.005 |  |  |
| **Q13510** | Acid ceramidase (ASAH1) | ↑1.88 | 0.001 | Y | [33] |
| Q9BQ51 | Programmed cell death 1 ligand 2 | ↑1.86 | 0.004 | Y | [43, 44] |
| **P11279** | Lysosome-associated membrane glycoprotein 1 | ↑1.83 | 0.028 | Y | [12] |
| **Q13228** | Selenium-binding protein 1 | ↑1.72 | 0.035 | Y | [45] |
| **P42785** | Lysosomal Pro-X carboxypeptidase | ↑1.63 | 0.012 | Y |  |
| P08236 | Beta-glucuronidase (GUSB) | ↑1.62 | 0.022 | Y | [32] |
| **P15144** | Aminopeptidase N | ↑1.59 | 0.043 |  | [39] |
| P01833 | Polymeric immunoglobulin receptor (pIgR) | ↑1.54 | 0.043 |  | [30] |
| **Q96PD5** | N-acetylmuramoyl-L-alanine amidase | ↓1.50 | 0.050 |  |  |
| P07339 | Cathepsin D (CTSD) | ↑1.50 | 0.050 | Y | [31] |

**Note:** Differential urinary proteins from both proteome analyses are in bold. <sup>a</sup> Change trends of proteins after tumor resection;
<sup>b</sup> protein expression in glioma tissues based on immunohistochemistry; <sup>c</sup> Proteins associated with gliomas in previous studies

**Figure 2.**
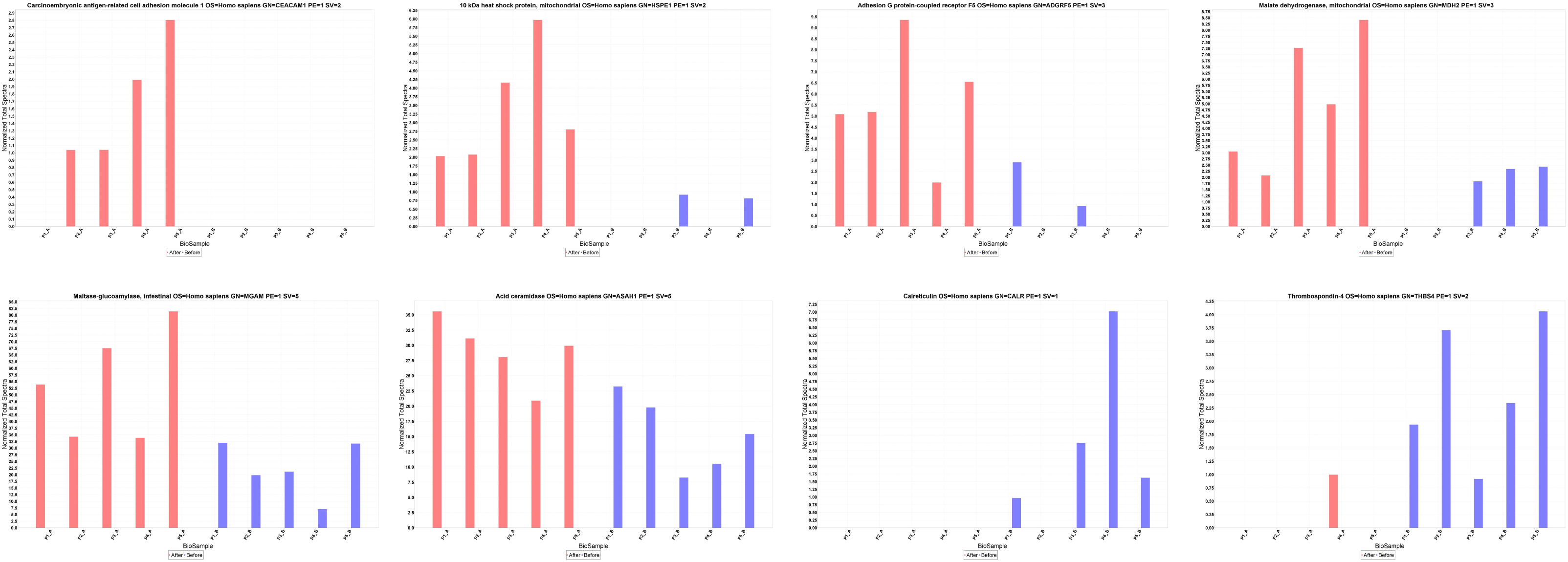
Several significantly altered urinary proteins identified by LC-MS/MS before and after tumor resection.

After reviewing the Human Protein Atlas Database available from www.proteinatlas.org, we discovered that 20 of these proteins are detectable in human glioma tissues using antibodies. These proteins might be secreted by glioma tissues and released into urine.

### 3. Functional analysis of differential urinary proteins

Functional annotation of the differential proteins was performed using DAVID, and they were classified as biological process, molecular function, molecular components and pathways. In the biological process category, regulation of tissue remodeling, autophagy, negative regulation of gene expression, positive regulation of natural killer cell-mediated cytotoxicity and angiogenesis were overrepresented (**Figure 3**). In the cellular component category, the majority of the identified proteins were extracellular exosome, lysosome, extracellular space and membrane proteins. In the molecular function category, peptide binding, integrin binding and metalloaminopeptidase activity were overrepresented. To identify the major biological pathways represented in the differential urinary proteins, KEGG was used for canonical pathway enrichment analysis. The pathways for the lysosome, the renin-angiotensin system, the phagosome and folate biosynthesis were significantly enriched.

**Figure 3.**
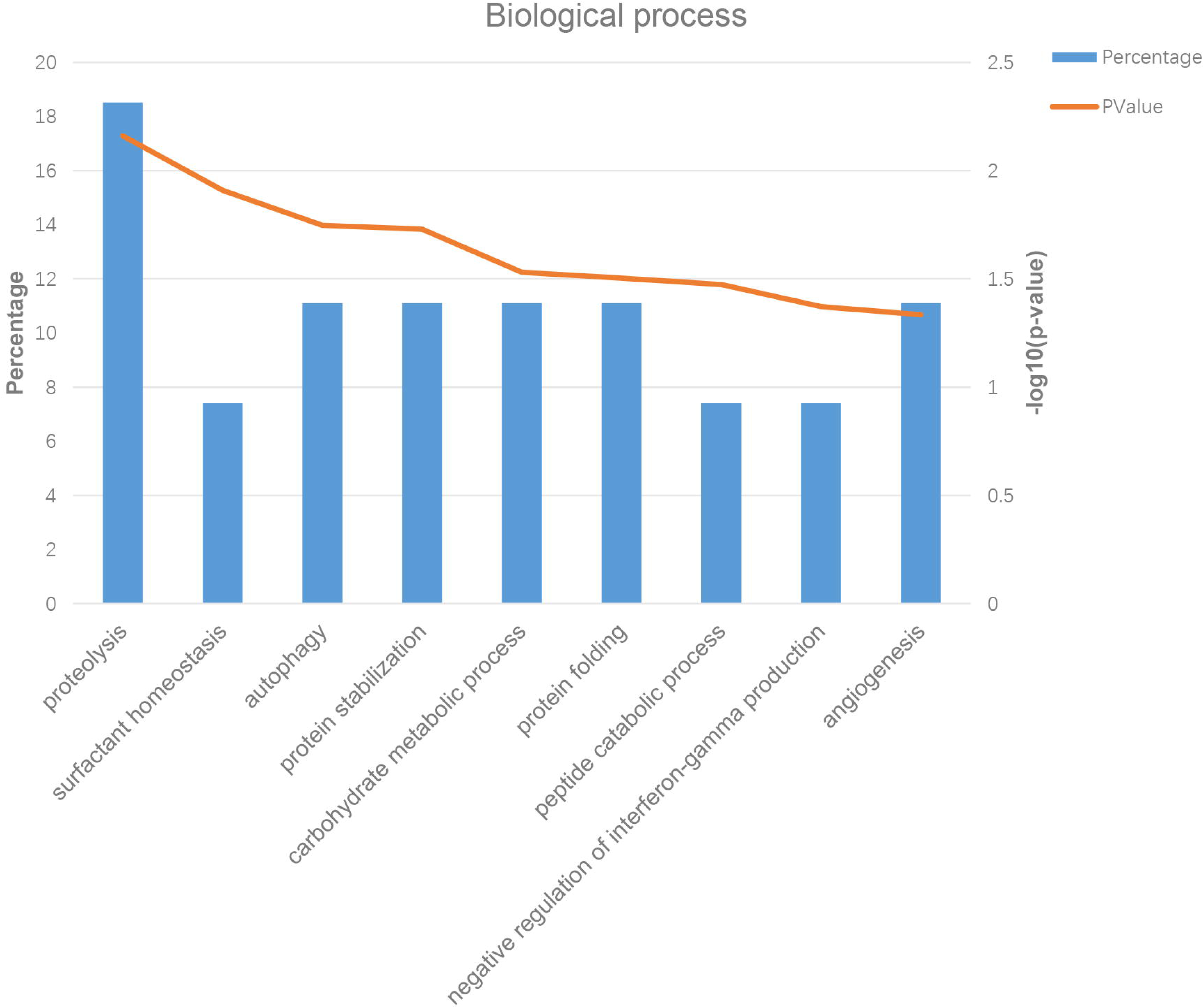
Functional annotation of differential urinary proteins before and after tumor resection.

Additionally, a total of 10 proteins were related to neurological diseases, including annexin A7, carboxypeptidase, vitellogenic-like, cathepsin D, crystallin alpha B, gamma-glutamyl hydrolase, lysosomal-associated membrane protein 1, nectin cell adhesion molecule 2, programmed cell death 1 ligand 2, quinoid dihydropteridine reductase, and thrombospondin 4.

### 4. Validation of differential urinary proteins

To identify more urinary proteins and validate altered protein level, the ten peptide samples were reanalyzed with an extended elution time of 90 min. A total of 1,652 urinary proteins were identified in this proteomic analysis (**Table S2**). Of the 27 proteins identified above, 17 proteins were verified to have significantly different levels in this analysis (**Table 2**).

## Discussion

In this study, a comparative proteomic analysis of urine samples from five glioma patients before and after tumor resection was performed using label-free proteome quantification. After brain tumor removal, the levels of 27 urinary proteins was significantly changed. Among these proteins, the abundance of 23 proteins was significantly increased after tumor removal, and four proteins were significantly decreased.

Of the differential urinary proteins, several have been previously associated with gliomas in previous studies, such as CEACAM1, ANXA7, CALR, CRYAB, CD276 antigen, pIgR, cathepsin D, beta-glucuronidase, and acid ceramidase (**Table 2**). For example, CEACAM1 controls matrix metalloproteinase-9 (MMP-9) secretion by neutrophils [19], and MMP-9 was reported to be a candidate brain cancer biomarker [2]. CEACAM1 silencing inhibits cell proliferation and promotes apoptosis in human glioma cells, suggesting that CEACAM1 is a potential glioma therapy target [20]. ANXA7 is a tumor suppressor gene. ANXA7 protein expression decreased in glioma tissues, and ANXA7 degradation might contribute to glioma progression [21]. Furthermore, loss of ANXA7 is associated with glioblastoma patient prognosis, and its expression was a strong predictor of patient outcome [22, 23]. CALR has also been implicated in cancer. Lower CALR levels are observed in glioma tissues than in normal brain tissues, and its expression is correlated with glioma tumor grade and patient survival [24]. Alpha-crystallin B chain (CRYAB) is an anti-apoptotic protein [25]. In glioblastoma multiforme (GBM), CRYAB levels were reportedly elevated [26], and CRYAB knockdown increased invasiveness in glioma cells [27]. CD276 antigen has been reported to be upregulated in high-grade glioma [28]. Its expression correlates with malignancy grade in gliomas and with poor patient survival [29]. Expression of polymeric immunoglobulin receptor (pIgR) was identified as a novel predictor of poor glioma patient prognosis after surgical resection [30]. Additionally, cathepsin D was identified as an important protein related to glioma invasion [31]. β-glucuronidase is an enzyme found in the necrotic area of GBM [32]. Acid ceramidase is a novel drug target for adjuvant pediatric brain tumor therapies [33]. Collectively, these urinary proteins might serve as noninvasive biomarker candidates to detect gliomas. Because these proteins were identified by comparing glioma patient samples before and after tumor resection, these proteins have potential clinical applications for diagnosing the presence of cancer and to monitor tumor recurrence after treatment in glioma patients.

Autophagy is involved in tumorigenesis, and a blockade has been proposed as an alternative therapeutic option. Autophagy is reportedly enhanced in astrocytomas compared to that in normal brain tissue [34]. In our study, autophagy was a significantly enriched biological process according to bioinformatics analyses, and four candidate proteins were involved in this process, including cathepsin D, lysosome-associated membrane glycoprotein 1, lysosome-associated membrane glycoprotein 2 and annexin A7.

Gliomas are characterized by abundant capillary angiogenesis [35], and antiangiogenic therapy is a promising approach to treat malignant gliomas [36]. In our study, several proteins were identified that are relevant to angiogenesis, such as CEACAM1 [37], CD276 [38], aminopeptidase N [39] and thrombospondin-4 [40]. Interestingly, thrombospondin-1 is involved in glioma pathogenesis and is a potential therapeutic target [41], but whether thrombospondin-4 plays a similar role requires further study.

Urine has been largely ignored during biomarker discovery for brain diseases compared to other bodily fluids, such as cerebrospinal fluid (CSF) and blood. However, few urinary-based biomarker studies examining brain diseases provided evidence that urine is a good source for relevant biomarkers [2, 42]. In this study, although a small number of clinical samples were analyzed, we identified several candidate biomarker proteins previously associated with gliomas in clinical studies. These proteins are likely to be validated in a large-scale urine sample analysis of glioma patients. As a preliminary study, our results show great potential for urine biomarker development to monitor gliomas or other brain cancers. However, a larger number of clinical urine samples from patients are needed to verify the specific proteins and protein patterns with clinical applications as noninvasive biomarkers for disease diagnosis and/or monitoring of tumor recurrence.

## Conclusion

Using label-free proteome quantification, candidate urinary biomarkers were identified in glioma patients with the potential to detect the disease and monitor tumor recurrence after treatment. The results suggested that inclusion of a self-control sample before tumor resection is an effective way to detect differential proteins related to the tumor, even in a small number of urine samples. Our findings may help future biomarker studies with clinical samples.

## Acknowledgement

This work was supported by National Key Research and Development Program of China (2016YFC1306300), Beijing Natural Science Foundation (7173264, 7172076), the Fundamental Research Funds for the Central Universities (2015KJJCB21), Beijing cooperative construction project (110651103) and Beijing Normal University (11100704).

## Supporting information

**Table S1**. Proteome profiling of glioma patient urine samples.

**Table S2**. Identification and quantitation details for the second urinary proteome analysis.

## References

[1] Ostrom, Q. T., Bauchet, L., Davis, F. G., Deltour, I., et al., The epidemiology of glioma in adults: a “state of the science” review. Neuro Oncol 2014, 16, 896–913.

[2] Smith, E. R., Zurakowski, D., Saad, A., Scott, R. M., Moses, M. A., Urinary biomarkers predict brain tumor presence and response to therapy. Clin Cancer Res 2008, 14, 2378–2386.

[3] Gao, Y., Urine-an untapped goldmine for biomarker discovery? Sci China Life Sci 2013, 56, 1145–1146.

[4] Gao, Y., Roadmap to the Urine Biomarker Era. MOJ Proteomics & Bioinformatics 2014, 1.

[5] Wang, Y., Chen, Y., Zhang, Y., Wu, S., et al., Differential ConA-enriched urinary proteome in rat experimental glomerular diseases. Biochem Biophys Res Commun 2008, 371, 385–390.

[6] Zhao, M., Li, M., Li, X., Shao, C., et al., Dynamic changes of urinary proteins in a focal segmental glomerulosclerosis rat model. Proteome Sci 2014, 12, 42.

[7] Yuan, Y., Zhang, F., Wu, J., Shao, C., Gao, Y., Urinary candidate biomarker discovery in a rat unilateral ureteral obstruction model. Sci Rep 2015, 5, 9314.

[8] Yin, W., Qin, W., Gao, Y., Urine glucose levels are disordered before blood glucose level increase was observed in Zucker diabetic fatty rats. Sci China Life Sci 2017.

[9] Wu, J., Guo, Z., Gao, Y., Dynamic changes of urine proteome in a Walker 256 tumor-bearing rats. Cancer Medicine 2017.

[10] Wu, J., Li, X., Gao, Y., Early detection in urinary proteome for the effective early treatment of bleomycin-induced pulmonary fibrosis in a rat model. 2017.

[11] Ni, Y., Zhang, F., An, M., Gao, Y., [Changes of urinary proteins in a bacterial meningitis rat model]. Sheng Wu Gong Cheng Xue Bao 2017, 33, 1145–1157.

[12] Ni, Y., Zhang, F., An, M., Yin, W., Gao, Y., Early candidate biomarkers found from urine of astrocytoma rat before changes in MRI. bioRxiv 2017, 117333.

[13] Zhang, F., Ni, Y., Yuan, Y., Yin, W., Gao, Y., Early urinary candidate biomarker discovery in a rat thioacetamide-induced liver fibrosis model. bioRxiv 2017, 125120.

[14] Zhao, M., Wu, J., Li, X., Gao, Y., Early urinary candidate biomarkers in a rat model of experimental autoimmune encephalomyelitis. 2017.

[15] Zhao, M., Wu, J., Li, X., Gao, Y., Urinary candidate biomarkers in an experimental autoimmune myocarditis rat model. 2017.

[16] Wu, J., Gao, Y., Physiological conditions can be reflected in human urine proteome and metabolome. Expert Rev Proteomics 2015, 12, 623–636.

[17] Old, W. M., Meyer-Arendt, K., Aveline-Wolf, L., Pierce, K. G., et al., Comparison of label-free methods for quantifying human proteins by shotgun proteomics. Mol Cell Proteomics 2005, 4, 1487–1502.

[18] Schmidt, C., Gronborg, M., Deckert, J., Bessonov, S., et al., Mass spectrometry-based relative quantification of proteins in precatalytic and catalytically active spliceosomes by metabolic labeling (SILAC), chemical labeling (iTRAQ), and label-free spectral count. RNA 2014, 20, 406–420.

[19] Ludewig, P., Sedlacik, J., Gelderblom, M., Bernreuther, C., et al., Carcinoembryonic antigen-related cell adhesion molecule 1 inhibits MMP-9-mediated blood-brain-barrier breakdown in a mouse model for ischemic stroke. Circ Res 2013, 113, 1013–1022.

[20] Xu, G., Li, W., Zhang, P., Ding, Z., et al., [Silencing of carcinoembryonic antigen-related cell adhesion molecule 1 inhibits proliferation and induces apoptosis in human glioma SHG44 cells]. Xi Bao Yu Fen Zi Mian Yi Xue Za Zhi 2015, 31, 23–26, 31.

[21] Pan, S. J., Zhan, S. K., Ji, W. Z., Pan, Y. X., et al., Ubiquitin-protein ligase E3C promotes glioma progression by mediating the ubiquitination and degrading of Annexin A7. Sci Rep 2015, 5, 11066.

[22] Ferrarese, R., Harsh, G. R. t., Yadav, A. K., Bug, E., et al., Lineage-specific splicing of a brain-enriched alternative exon promotes glioblastoma progression. J Clin Invest 2014, 124, 2861–2876.

[23] Hung, K. S., Howng, S. L., Prognostic significance of annexin VII expression in glioblastomas multiforme in humans. J Neurosurg 2003, 99, 886–892.

[24] Gao, H., Yu, B., Yan, Y., Shen, J., et al., Correlation of expression levels of ANXA2, PGAM1, and CALR with glioma grade and prognosis. J Neurosurg 2013, 118, 846–853.

[25] Kore, R. A., Abraham, E. C., Phosphorylation negatively regulates exosome mediated secretion of cryAB in glioma cells. Biochim Biophys Acta 2016, 1863, 368–377.

[26] Kore, R. A., Abraham, E. C., Inflammatory cytokines, interleukin-1 beta and tumor necrosis factor-alpha, upregulated in glioblastoma multiforme, raise the levels of CRYAB in exosomes secreted by U373 glioma cells. Biochem Biophys Res Commun 2014, 453, 326–331.

[27] Shimizu, M., Tanaka, M., Atomi, Y., Small Heat Shock Protein alphaB-Crystallin Controls Shape and Adhesion of Glioma and Myoblast Cells in the Absence of Stress. PLoS One 2016, 11, e0168136.

[28] Zhou, Z., Luther, N., Ibrahim, G. M., Hawkins, C., et al., B7-H3, a potential therapeutic target, is expressed in diffuse intrinsic pontine glioma. J Neurooncol 2013, 111, 257–264.

[29] Lemke, D., Pfenning, P. N., Sahm, F., Klein, A. C., et al., Costimulatory protein 4IgB7H3 drives the malignant phenotype of glioblastoma by mediating immune escape and invasiveness. Clin Cancer Res 2012, 18, 105–117.

[30] Niu, H., Wang, K., Wang, Y., Polymeric immunoglobulin receptor expression is predictive of poor prognosis in glioma patients. Int J Clin Exp Med 2014, 7, 2185–2190.

[31] Pei, J., Moon, K. S., Pan, S., Lee, K. H., et al., Proteomic Analysis between U87MG and U343MG-A Cell Lines: Searching for Candidate Proteins for Glioma Invasion. Brain Tumor Res Treat 2014, 2, 22–28.

[32] Bensalma, S., Chadeneau, C., Legigan, T., Renoux, B., et al., Evaluation of cytotoxic properties of a cyclopamine glucuronide prodrug in rat glioblastoma cells and tumors. J Mol Neurosci 2015, 55, 51–61.

[33] Doan, N. B., Nguyen, H. S., Montoure, A., Al-Gizawiy, M. M., et al., Acid ceramidase is a novel drug target for pediatric brain tumors. Oncotarget 2017, 8, 24753–24761.

[34] Jennewein, L., Ronellenfitsch, M. W., Antonietti, P., Ilina, E. I., et al., Diagnostic and clinical relevance of the autophago-lysosomal network in human gliomas. Oncotarget 2016, 7, 20016–20032.

[35] Takano, S., Yamashita, T., Ohneda, O., Molecular therapeutic targets for glioma angiogenesis. J Oncol 2010, 2010, 351908.

[36] Takano, S., Kamiyama, H., Tsuboi, K., Matsumura, A., Angiogenesis and antiangiogenic therapy for malignant gliomas. Brain Tumor Pathol 2004, 21, 69–73.

[37] Muller, M. M., Singer, B. B., Klaile, E., Obrink, B., Lucka, L., Transmembrane CEACAM1 affects integrin-dependent signaling and regulates extracellular matrix protein-specific morphology and migration of endothelial cells. Blood 2005, 105, 3925–3934.

[38] Seaman, S., Stevens, J., Yang, M. Y., Logsdon, D., et al., Genes that distinguish physiological and pathological angiogenesis. Cancer Cell 2007, 11, 539–554.

[39] Mayas, M. D., Ramirez-Exposito, M. J., Carrera, M. P., Cobo, M., Martinez-Martos, J. M., Renin-angiotensin system-regulating aminopeptidases in tumor growth of rat C6 gliomas implanted at the subcutaneous region. Anticancer Res 2012, 32, 3675–3682.

[40] Muppala, S., Xiao, R., Krukovets, I., Verbovetsky, D., et al., Thrombospondin-4 mediates TGF-beta-induced angiogenesis. Oncogene 2017, 36, 5189–5198.

[41] Ma, Y., Qu, B., Xia, X., Yang, L., et al., Glioma-derived thrombospondin-1 modulates cd14+ cell tolerogenic properties. Cancer Invest 2015, 33, 152–157.

[42] An, M., Gao, Y., Urinary Biomarkers of Brain Diseases. Genomics Proteomics Bioinformatics 2015, 13, 345–354.

[43] Miyazaki, T., Ishikawa, E., Matsuda, M., Akutsu, H., et al., Assessment of PD-1 positive cells on initial and secondary resected tumor specimens of newly diagnosed glioblastoma and its implications on patient outcome. J Neurooncol 2017, 133, 277–285.

[44] Wang, Z., Zhang, C., Liu, X., Wang, Z., et al., Molecular and clinical characterization of PD-L1 expression at transcriptional level via 976 samples of brain glioma. Oncoimmunology 2016, 5, e1196310.

[45] Rajaraman, P., Brenner, A. V., Butler, M. A., Wang, S. S., et al., Common variation in genes related to innate immunity and risk of adult glioma. Cancer Epidemiol Biomarkers Prev 2009, 18, 1651–1658.

